# Differential Associations of Microglial Inflammation on LATE-NC and Tangle-Related Hippocampal Atrophy

**DOI:** 10.64898/2026.08.24.744255

**Authors:** A Kapasi, L Yu, SE Leurgans, EY Chen, S Agrawal, LL Barnes, DA Bennett, K Arfanakis, JA Schneider

**Affiliations:** Rush Alzheimer’s Disease Center, Rush University Medical Center, Chicago, IL, USA; Department of Pathology, Rush University Medical Center, Chicago, IL, USA; Department of Neurological Sciences, Rush University Medical Center, Chicago, IL, USA; Rush University Medical Center, Department of Diagnostic Radiology and Nuclear Medicine, Chicago, IL; Illinois Institute of Technology, Department of Biomedical Engineering, Chicago, IL

**Keywords:** Microglia, LATE-NC, AD, hippocampal volume, ex-vivo MRI, digital pathology

## Abstract

**BACKGROUND:** Accumulations of AD and LATE-NC both contribute to changes in hippocampal volume, possibly via distinct and/or overlapping mechanisms. Microglia-driven inflammation is a shared pathway associated with both AD and LATE-NC. However, the extent to which microglia inflammation is associated with hippocampal volume is less understood.

**OBJECTIVE:** Examine the relationship between AD and LATE-NC with hippocampal volume in persons with differing levels of microglia inflammation.

**METHODS:** Cerebral hemispheres from 441 older adults who came to autopsy were studied. All hemispheres underwent ex-vivo MRI and detailed neuropathologic examination for neurodegenerative and cerebrovascular pathologies. Microglia were quantified in the hippocampal CA1/subiculum region using machine learning-based classifiers trained on digitized CR3-43-stained images via the HALO digital pathology platform. First, linear regression models examined the association of microglia with hippocampal volume, adjusting for demographics, postmortem interval (PMI), and common age-related pathologies. Second, linear regression models were employed to examine whether microglia density modified associations of β-amyloid, tangle, or LATE-NC on hippocampal volume.

**RESULTS:** Participants had a mean age of 90 years at death with 75% being women. Intermediate or high likelihood ADNC was present in 64% and LATE-NC (stage 2/3) was present in 52%. In linear regression models, adjusting for demographics and PMI, higher microglia density was associated with a lower hippocampal volume to hemisphere ratio (estimate = −0.021 SE=0.01, p=0.002); however, after adjusting for common age-related pathologies the association was attenuated (p=0.70). β-amyloid, tangles, and LATE-NC remained independently associated with a lower hippocampal volume. The association of LATE-NC with hippocampal volume was stronger in brains with greater microglia burden (estimate for the interaction term = −0.016; SE=0.01, p=0.002). No interactions were seen between β-amyloid or tangles with microglia on hippocampal volume. In stratified analyses, microglial density modified the association between LATE-NC and hippocampal volume, independent of AD neuropathologic status.

**CONCLUSION:** Microglia-driven inflammation strengthens the association of LATE-NC, but not AD pathology, on hippocampal volume loss. These findings emphasize the importance of inflammatory pathways [when interpreting MRI-based neurodegeneration markers] in aging and mixed pathology.

## INTRODUCTION

Hippocampal atrophy represents a well-characterized neuroimaging and neuropathological marker of cognitive decline associated with aging and neurodegeneration. Both Alzheimer’s disease (AD) pathology and limbic-predominant age-related TDP-43 encephalopathy neuropathologic change (LATE-NC) have been identified as major pathologies contributing to hippocampal atrophy^1–4^. Although these proteinopathies frequently co-occur within the aging brain ^5,6^, evidence is accumulating that they may drive hippocampal neurodegeneration through distinct, yet potentially convergent, pathogenic mechanisms.

Microglia-mediated neuroinflammation has emerged as a central convergent mechanism linking multiple neurodegenerative processes to neuronal and synaptic loss. Upon activation, microglia secrete proinflammatory cytokines, modulate the local cellular milieu, and potentiate neurotoxic cascades, thereby amplifying the downstream consequences of neurodegenerative processes ^7–10^. Several notable studies have examined the relationship between microglia inflammation and regional brain atrophy. In-vivo PET imaging studies have shown that greater microglial activation is associated with greater cortical thinning, particularly in parietal, occipital, and cingulate regions ^11^. Complementary histopathological evidence from cases with primary progressive aphasia and frontotemporal lobar degeneration (FTLD) indicate that white matter underlying atrophied cortical areas exhibit higher microglial burden compared to non-atrophied regions ^12^. Moreover, brain regions enriched in disease-associated genetic risk variants in microglia are often the same regions that exhibit brain atrophy in AD ^13^. However, few studies quantify the association of microglial inflammation in the hippocampus with hippocampal atrophy. Our prior work has shown that both LATE-NC and tau-tangle pathology are independently associated with regionally elevated microglia density within the hippocampus^14^; however, the extent to which microglia modulate the association of these proteinopathies on hippocampal atrophy remains poorly understood.

To address these questions, we examined 440 older persons with ex-vivo MRI, detailed neuropathologic assessments for AD and LATE-NC, and generated quantitative digital histopathology data for microglia density. Specifically, we first investigated whether microglial density in hippocampus is associated with hippocampal volume and secondly, whether microglial density modifies the relationship between AD or LATE-NC on hippocampal volume.

## METHODS

### Participants

Participants enrolled in the Religious Orders Study (ROS), Rush Memory and Aging Project (MAP), Clinical Core, or Minority Aging Research Study (MARS) ^15^. Eligibility for each study requires the absence of known dementia, and all participants agree to annual clinical evaluation and interview. For ROS and MAP, participants agree to undergo brain donation, while Clinical Core and MARS brain donation is optional. Participants signed an informed consent, and an Anatomical Gift Act for brain donation. Each study was approved by Rush University Medical Center institutional review board ^16^.

### Clinical Assessment

All participants underwent a standardized annual clinical evaluation that included a medical history, neurologic examination, and a battery of 19 cognitive tests. Cognitive impairment was initially assessed by a neuropsychologist who reviewed clinical data while blinded to participant demographic characteristics. A clinician subsequently reviewed all available clinical information, examined the participant, and rendered a final clinical diagnosis. To determine cognitive status proximate to death, a clinician masked to postmortem findings reviewed longitudinal clinical data from all study visits and assigned a final diagnosis ^17^.

Clinical diagnoses were assigned according to established consensus criteria ^18^, Dementia required cognitive decline from a prior level of functioning with impairment in multiple cognitive domains. Participants with cognitive impairment who did not meet criteria for dementia were classified as having mild cognitive impairment (MCI), and all others were classified as having no cognitive impairment (NCI).

### Ex Vivo MRI Acquisition and Hippocampal Volumes Measures

At autopsy, the hemisphere exhibiting the greatest pathology burden or least mechanical damage was fixed in 4% phosphate-buffered formaldehyde at 4°C. Approximately 30 days postmortem, the intact hemisphere underwent ex vivo MRI with the medial surface oriented downward, as previously described ^19^. MRI data were acquired on 3T scanners using a multi-echo spin-echo sequence (voxel size = 0.63 × 0.63 × 1.5 mm³; echo times = 11–82.5 ms; repetition time ≈ 4000 ms; sagittal acquisition; total scan time ≈ 30 minutes).¹

The hippocampus and hemisphere were segmented using an in-house multi-atlas segmentation pipeline. Prior work has demonstrated that ex vivo regional brain volumes remain stable for up to six months postmortem and are linearly associated with corresponding in vivo MRI measurements, supporting the validity of ex vivo MRI as a measure of antemortem brain structure ^1^. Hippocampal volume was calculated by multiplying the number of segmented hippocampal voxels by voxel volume. For analyses, hippocampal volume was expressed as the ratio of hippocampal-to-hemisphere volume × 10³, consistent with prior studies ^20^. This measure is referred to as “hippocampal volume” throughout the manuscript.

### Diagnostic Neuropathology

The median post-mortem interval was 7.92 hours (interquartile range: 6.17 - 12.50). Brain removal and processing followed a standardized protocol. Briefly, the fixed intact hemisphere designated for ex vivo MRI was sectioned into 1-cm-thick coronal slabs and stored in cryoprotectant solution. Fixed tissue samples from a standard set of at least 12 cortical, subcortical, brainstem, and cerebellar regions were collected for diagnostic evaluation, microscopy, and digital pathology analyses. Tissue blocks were dehydrated, embedded in paraffin, sectioned at 6 μm, and stained using established protocols. Neuropathologic assessments were performed by investigators blinded to participant demographic and clinical information

#### Neurodegenerative Pathologies – Pathologic Diagnosis of AD

Alzheimer’s disease (AD) neuropathologic diagnosis was determined according to NIA-AA criteria ^21^. Modified Braak stage and CERAD scores were derived from manual assessments of neurofibrillary tangles and neuritic plaques using a modified Bielschowsky stain across five brain regions. Thal amyloid phase was assigned based on the regional distribution of β-amyloid pathology across seven brain regions, including the neocortex, hippocampus, basal ganglia, midbrain, and cerebellum.

#### β-amyloid load

Quantitative measures for β-Amyloid load are based on historical and/or digital data from 8 brain regions including middle and superior frontal, inferior temporal, cingulate gyrus, inferior parietal, calcarine, and entorhinal cortices, and hippocampus. Cortical and hippocampal regions of interest were manually outlined (excluding meninges). Digital image analysis for Aß pathology was quantified using the Aperio Image Analysis Toolbox Positive Pixel Count algorithm. Digital measures for β-amyloid are highly correlated with historical data and have been harmonized, as previously described ^22^.

#### PHF tau-tangles density

Quantitative measures for tangles density are based on historical and/or digital data from 8 brain regions including middle and superior frontal, inferior temporal, cingulate gyrus, inferior parietal, calcarine, and entorhinal cortices, and hippocampus. Cortical and hippocampal regions of interest were manually outlined. Digital image analysis for tau tangle pathology was quantified using the Aperio Image Analysis Toolbox Nuclear algorithm. Digital measures for tau tangles are highly correlated with historical data and have been harmonized, as previously described ^22^.

#### LATE-NC

TDP 43 pathology was assessed by immunohistochemistry using a monoclonal antibody to phosphorylated TDP 43 (pS409/410; 1:100). TDP 43 was evaluated in 8 brain regions: amygdala, hippocampus CA1-subiculum, dentate gyrus, entorhinal cortex, anterior temporal pole, middle temporal gyrus, midfrontal gyrus, and orbitofrontal cortex ^23^.

#### Lewy body pathology

Lewy bodies were evaluated using a phosphorylated α synuclein antibody (Wako; 1:20 000) in 7 brain regions, as described previously ^24^.

#### Cerebrovascular Pathologies – Infarcts

Macroscopic infarcts were identified on gross examination and characterized by location, size, and age, with infarct age confirmed histologically as acute, subacute, or chronic. Microscopic infarcts were identified by histologic examination and similarly classified. Only chronic macroscopic and microscopic infarcts were included in analyses ^25^.

#### Arteriolosclerosis

Basal ganglia arteriolosclerosis was evaluated on hematoxylin and eosin– stained sections and graded using a semi-quantitative 4-point scale (0 = none, 1 = mild, 2 = moderate, and 3 = severe). Ratings were based on the severity of arteriolar wall changes, including intimal degeneration, smooth muscle cell loss, concentric hyaline thickening, and narrowing of the vascular lumen ^26^.

#### Atherosclerosis

Intracranial atherosclerosis was evaluated semi-quantitatively in the circle of Willis and major cerebral arteries, including the vertebral, basilar, posterior cerebral, middle cerebral, and anterior cerebral arteries and their proximal branches. Severity ratings incorporated plaque burden, extent of vessel involvement, and degree of luminal occlusion ^20^.

### Digital Scanning

Hippocampal slides stained with CR3-43 (Abcam; 1:100) were digitized at 20x magnification to obtain whole slide images using the Leica Aperio AT2 console. Quality control was performed for each whole slide image to ensure suitability for digital image analysis. Images were carefully inspected for staining and/or tissue sectioning-related artifacts that may impact downstream data quality ^22^.

### Quantification of Hippocampal (CA1/subiculum) Microglia

#### Quantification via machine learning-based classifiers

Using the HALO AI platform (Indica Labs), we employed an untrained version of the nuclei segmentation classifier to first detect total microglial cells. The classifier was trained on manually annotated digital images to distinguish microglial cells from background. This training dataset included 750 annotations for microglial cell segmentation and 35 annotations for background. Model performance was refined through iterative cross-validation and visual inspection of detection overlays in HALO AI. Misclassifications were corrected by adjusting training parameters and refining annotations. Because our prior work has shown that CR3-43-stained microglia cells show various morphologies (ramified, hypertrophic, and ameboid) ^14^, we integrated the nuclei segmentation classifier for total microglia with the HALO AI Object Phenotype module. This allowed quantification of cell phenotypes into two separate groups: Stage 1 and 1d which included long/short, ramified processes with cell bodies <14um, and Stage 2/3 which included cell bodies >14um with hypertrophic and/or ameboid morphology. The HALO AI Object Phenotype module provides a predictability confidence score ranging from 0.01 (low confidence) to 1.00 (high confidence) for each cell assigned a phenotype. To ensure high classification accuracy, only microglia stage 2/3 cells with predictability confidence scores of >0.95 were included when calculating density of stage 2/3 microglia (Figure 1).

**Figure 1.**
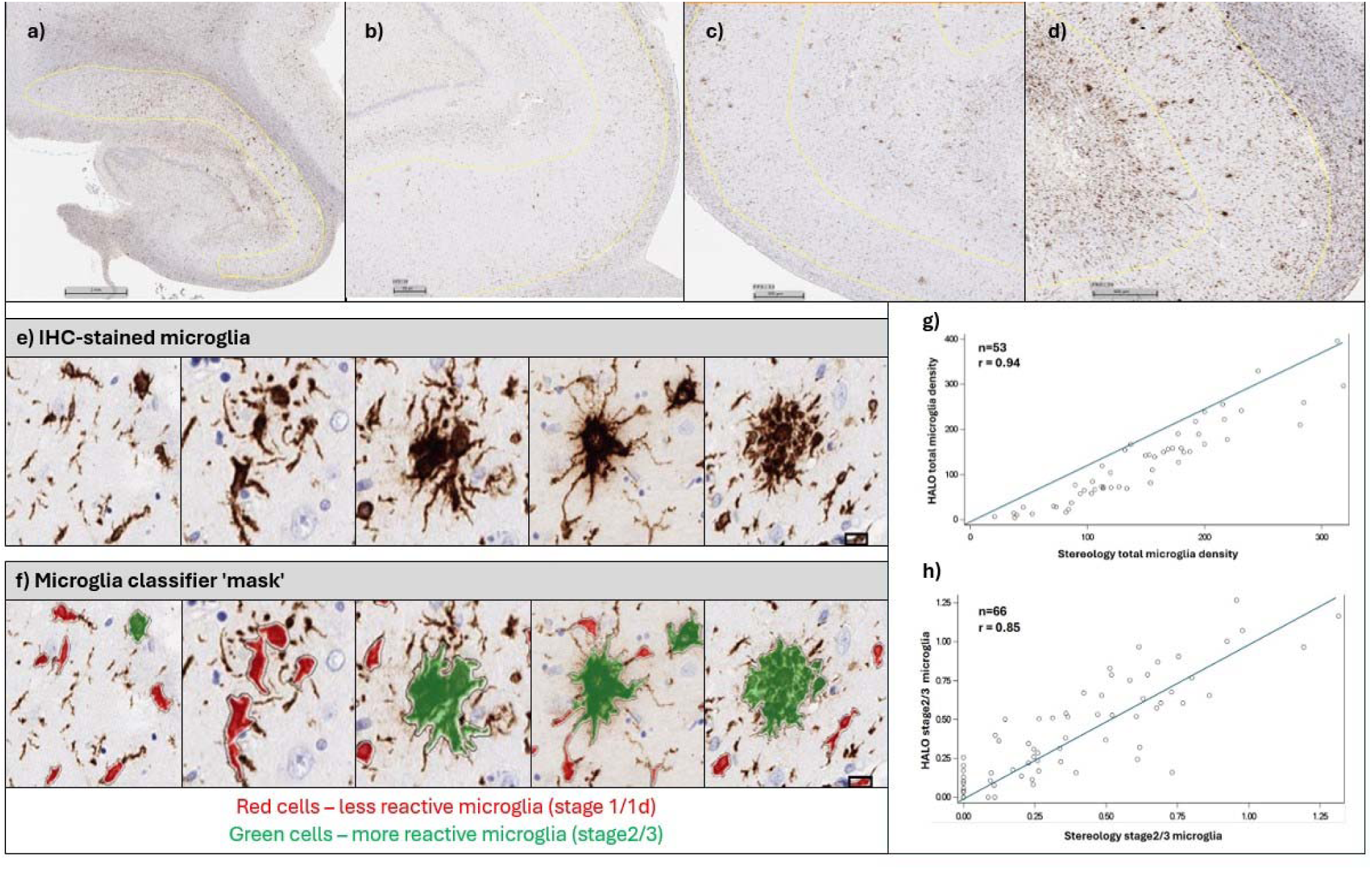
Machine learning-based pathology classifier for the quantification of microglia. Panel (a) demonstrates an example of a CA1-subiculum ROI outline at low magnification. Panels (b-d) demonstrate image examples of low, intermediate, and high density of hippocampal microglia. Panel (e) and (f) demonstrate images of microglial cells from CR3-43-stained sections and the microglia classifier ‘mask’. Panels (g) and (h) demonstrate the correlation between digital-derived and manual counting of total and stage2/3 microglia.

#### Quantification via manual stereology

Manual stereology counts were performed using the StereoInvestigator 8.0 software. Hippocampal CA1/Subiculum sub-regions were manually outlined, and counting parameters were set up to sample 15% of the region. Counts were performed by expert raters with extensive experience, and microglial cells were grouped based on each morphologic stage, as previously described ^14^. For analyses, 441 participants were included that had valid baseline, ex-vivo MRI data for hippocampal volume, and a complete neuropathologic diagnostic workup and digital quantification of microglia.

### Statistics

#### Validation of digital-derived data with manual stereology counts for microglia

In order to validate our digital classifier for microglia, we first examined spearman correlations between the digital-derived data with the manual stereology counts for microglia in a subset of cases (n=53 for total microglia and n=66 for Stage 2/3 microglia). Digital counts (without data transformation) showed a strong positive correlation with stereology for both total microglia (r_s_= 0.93, <0.001) and Stage 2/3 microglia (r_s_= 0.83, <0.001). These findings demonstrate high concordance between the digital and manual methods, supporting the validity of the digital classifier (Figure 1). The digital data for total hippocampal microglia showed a skew distribution, so analyses employed the square-root transformation. Like our prior work ^14^, the density measure for Stage 2/3 microglia, derived from only those stage 2/3 cells that have a predictability score >0.95 divided by the same area outlined, also exhibited a skewed distribution and required log transformation prior to analysis.

#### Analytic Approach

Spearman correlation (r_s_) was used to examine bivariate analyses of hippocampal microglia with age, education, β-amyloid, and tangles. One-way ANOVA tests were used to compare microglia across pathologic diagnoses. For description, we divided the continuous values of microglia density into tertiles to create low, intermediate, and high microglia burden groups. Linear regression models were used to examine the association of total microglia with hippocampal volume. The first model included the terms for demographics (age, sex, education), scanner, and postmortem intervals to fixation and to imaging with hippocampal volume as the outcome. The second model included nine additional pathology terms (cortical β-amyloid, tau tangles, LATE-NC, Lewy bodies, and cerebrovascular pathologies). We repeated analyses replacing total microglia density with stage2/3 microglia density. Because this model only showed that β-amyloid, tangles, and LATE-NC were significantly associated with hippocampal volume, we examined whether microglia burden influenced the association of these pathologies on hippocampal volume. To examine the interaction between β-amyloid and microglia, we fitted a linear regression model with hippocampal volume as the outcome and terms for β-amyloid and microglia, an interaction term between β-amyloid and microglia, and terms to adjust for demographics, scanner, PMI to fixation and imaging, and other brain pathologies. We repeated the analysis by replacing the β-amyloid term with tangles and separately with LATE-NC. In subsequent analyses, we fitted linear regression models with an interaction term between microglia and LATE-NC with hippocampal volume as the outcome, stratified by presence or absence of AD neuropathologic diagnosis.

All analyses were programmed in SAS/STAT, version 9.4 [SAS Institute Inc., Cary, NC]. Statistical significance was determined at a nominal α level of 0.05.

## RESULTS

Demographic, clinical, and neuropathologic characteristics of 441 older participants are presented in Table 1. On average, mean age at death was 90.1 years (SD=6.82) years and mean education was 16 years. The mean total microglia density was 97.92 with a range of 0.07 to 621.96. Stage 2/3 microglia had a mean density of 0.36 (SD=0.38). Total microglia were weakly correlated with lower hippocampal volume (r_s_= −0.149, p=0.002) and higher β-amyloid load (r_s_=0.232, p<0.001), tangles density (r_s_=0.385, p<0.001), LATE-NC stage {F(3,425) = 13.97, <0.001}, Lewy bodies {t(193) =-2.62, p=0.009}, arteriolosclerosis severity {F(3,425) = 2.65, *p*=0.048} and atherosclerosis severity {F(3,425) = 3.35, *p*=0.019}. As expected, cases with more dementia, a higher likelihood of AD neuropathologic change, and more neocortical LATE-NC, had higher levels of microglia density (Table 1).

**Table 1.** Characteristics of participants.

|  | All (n=441) | Low Microglia<br>Density<br>(n=147) | Intermediate<br>Microglia Density<br>(n=147) | High Microglia<br>Density<br>(n=147) |
| --- | --- | --- | --- | --- |
| **Demographics** |  |  |  |  |
| Mean age at death, years | 90.1 | 88.6 | 90.6 | 91.2 |
| Women, n (%) | 332 (75%) | 111 (76%) | 110 (75%) | 111 (76%) |
| Education, years | 15.6 | 15.5 | 15.9 | 15.4 |
| **Clinical dx, n (%)** |  |  |  |  |
| Normal Cognition | 155 (35%) | 76 (52%) | 51 (35%) | 28 (19%) |
| MCI | 94 (21%) | 29 (20%) | 37 (25%) | 28 (19%) |
| Dementia | 192 (44%) | 42 (29%) | 59 (40%) | 91 (62%) |
| **Ex-vivo MRI volumes, mL** |  |  |  |  |
| Hippocampal Volume | 3,051 | 3,167 | 3,096 | 2,891 |
| Hemisphere Volume | 363,352 | 371,178 | 366,576 | 352,301 |
| **Neuropathology** |  |  |  |  |
| ADNC, NIA-AA criteria |  |  |  |  |
| None | 43 (10%) | 13 (9%) | 16 (11%) | 14 (10%) |
| Low | 105 (24%) | 50 (34%) | 39 (27%) | 16 (11%) |
| Intermediate | 175 (40%) | 65 (44%) | 56 (38%) | 54 (37%) |
| High | 118 (27%) | 19 (13%) | 36 (24%) | 63 (43%) |
| β-amyloid load | 1.16 | 0.97 | 0.99 | 1.23 |
| Tangle density | 1.52 | 1.09 | 1.41 | 2.06 |
| LATE-NC Stage |  |  |  |  |
| None | 187 (42%) | 71 (48%) | 71 (48%) | 45 (31%) |
| Amygdala | 85 (19%) | 36 (24%) | 29 (20%) | 20 (14%) |
| Limbic | 48 (11%) | 15 (10%) | 18 (20%) | 15 (10%) |
| Neocortical | 121 (27%) | 25 (17%) | 29 (20%) | 67 (46%) |
| Lewy bodies | 128 (29%) | 37 (25%) | 41 (28%) | 50 (34%) |
| Macroscopic Infarcts | 146 (33%) | 46 (31%) | 49 (33%) | 51 (35%) |
| Microscopic Infarcts | 144 (33%) | 44 (30%) | 50 (34%) | 50 (34%) |
| Mod/Sev atherosclerosis | 72 (16%) | 11 (7%) | 31 (21%) | 30 (20%) |
| Mod/Sev arteriolosclerosis | 109 (24%) | 27 (18%) | 38 (26%) | 44 (30%) |

### Microglia and Hippocampal Volume

First, we examined the association of total microglia with hippocampal-to-hemisphere volume ratio in a linear regression model adjusted for demographics, scanner, and postmortem interval. In this model, total microglia were related to a lower hippocampal-to-hemisphere volume ratio. However, when neurodegenerative and cerebrovascular pathologies were included in the model, the association of total microglia density with hippocampal volume was attenuated, which suggests that microglia density is not independently associated with hippocampal atrophy above and beyond other coexisting age-related pathological processes. β-amyloid, tangle density, and LATE-NC were independently related to a lower hippocampal-to-hemisphere volume ratio (Table 2). We examined the associations of stage 2/3 microglia with hippocampal-to-hemisphere volume and similar associations were found (supplementary Table 1)

**Table 2.** Association of total microglia density with hippocampal to hemisphere volume ratio.

| Predictors | Hippocampal to Hemisphere Volume Ratio <sup>a</sup> | Hippocampal to Hemisphere Volume Ratio <sup>b</sup> |
| --- | --- | --- |
| Total Microglia | -0.372 (0.14,0.01) | 0.135 (0.15, 0.38) |
| β-Amyloid |  | -2.896 (1.22, 0.02) |
| Tangles |  | -1.706 (0.74, 0.02) |
| LATE-NC |  | -2.896 (0.55, <0.001) |
| Lewy bodies |  | 0.949 (1.45,0.51) |
| Atherosclerosis |  | -1.026 (1.10, 0.35) |
| Arteriolosclerosis |  | 0.613 (0.81,0.45) |
| Macroscopic infarcts |  | -3.072 (1.47,0.05) |
| Microscopic infarcts |  | 0.945 (1.45,0.52) |

<sup>a</sup> Linear regression model of hippocampal volume as the outcome on total microglia density, with terms to adjust for age at death, sex, education, PMI to fixation, PMI to imaging, and MRI scanner. In analyses, total microglia applied with sq-root transformation was used.
<sup>b</sup> Linear regression model of hippocampal volume as the outcome, included 9 terms of pathologies and terms to adjust for age at death, sex, education, PMI to fixation, PMI to imaging, and MRI scanner. In analyses, total microglia applied with sq-root transformation was used.

### Interaction of AD and LATE-NC with Microglia on Hippocampal Volume

Next, we examined whether microglia density modifies the association of β-amyloid, tangles, or LATE-NC with hippocampal volume. In linear regression models with an interaction term between β-amyloid and total microglia, as well as between tangles and total microglia, we observed no interaction on hippocampal-to-hemisphere volume ratio. However, these models also showed that regardless of microglia density both tangles and β-amyloid were associated with lower hippocampal volume; suggesting AD pathology may drive hippocampal change via other biologic mechanisms beyond microglia inflammation in the hippocampus. In contrast, we found an interaction between LATE-NC and total microglia, such that with each additional 1-unit of microglia density, LATE-NC was associated with an additional 0.019-units lower hippocampal volume (Table 3, Figure 2).

**Figure 2.**
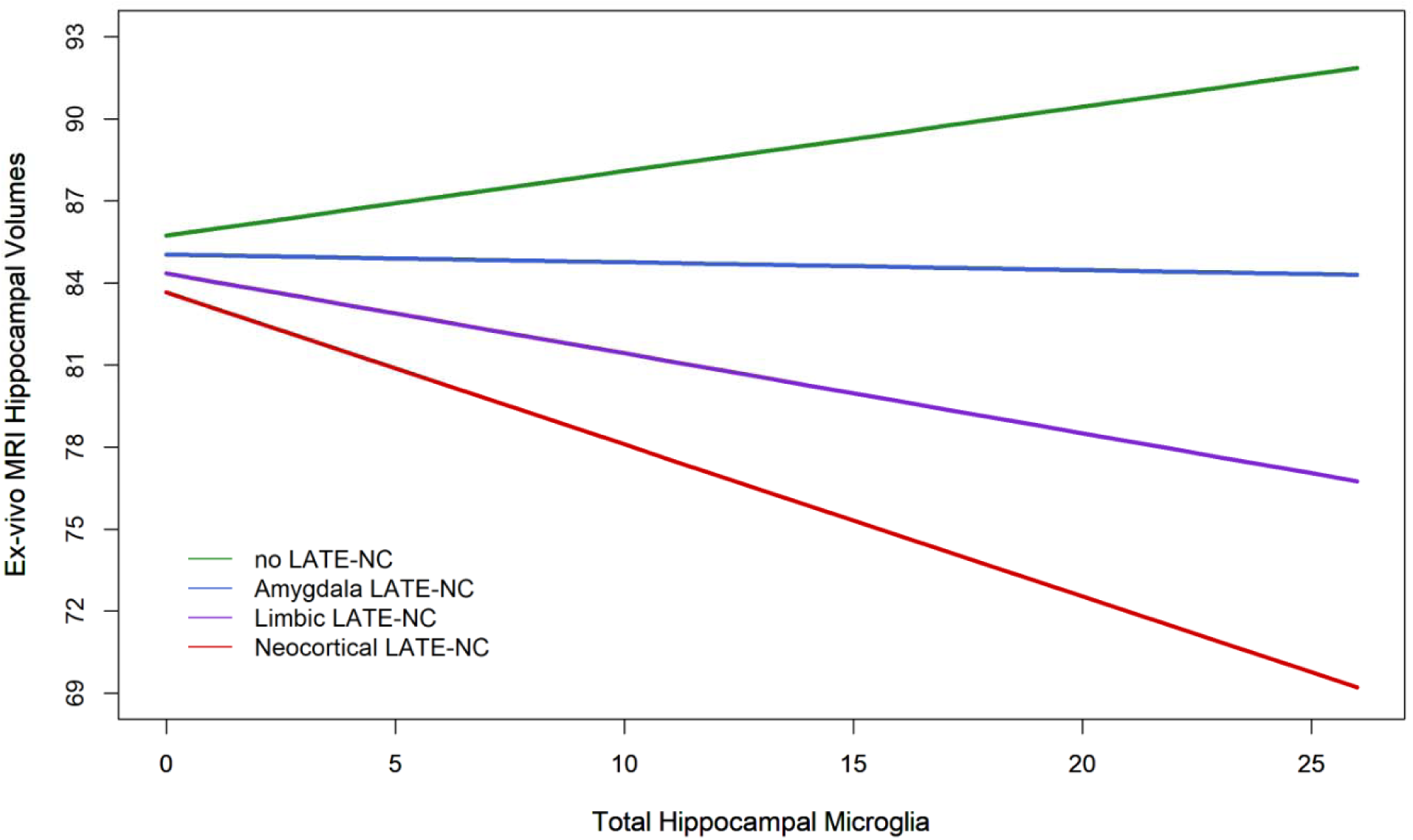
Estimated hippocampal volume versus level of microglia density by LATE-NC stage.

**Table 3.**
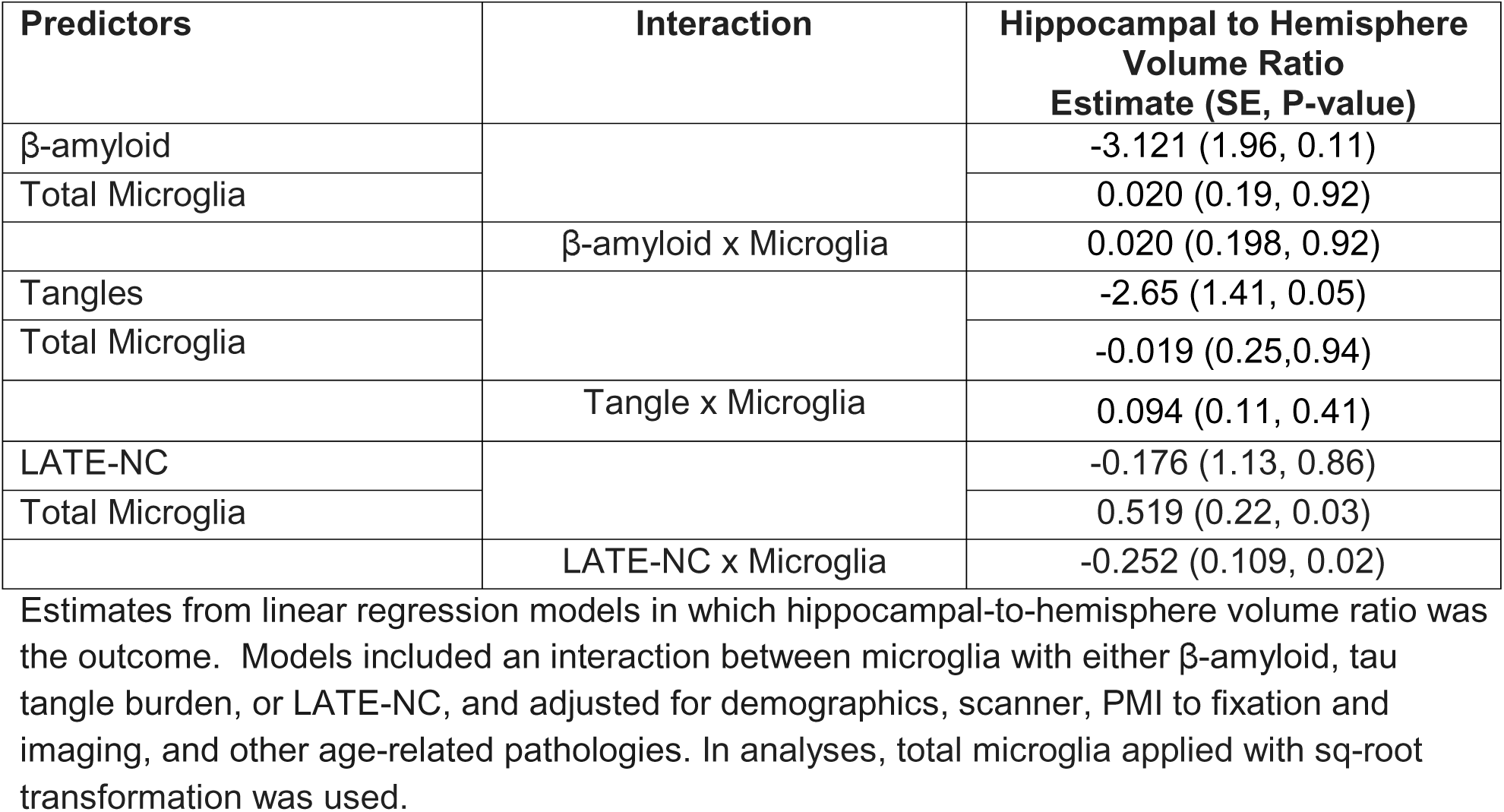
Interaction of AD pathology and LATE-NC with microglia density in relation to hippocampal volume.

In subsequent analyses, we examined whether the interaction between LATE-NC and total microglia on hippocampal volume stratified by AD neuropathologic diagnosis. We found that there was an interaction between LATE-NC and microglia on hippocampal volume among those with an AD neuropathologic diagnosis (n=293; Interaction estimate = −0.326, SE =0.138, p=0.018) and among those without an AD neuropathologic diagnosis (n=148; Interaction estimate= −0.390, SE=0.187, p=0.039). Together, these findings suggest that microglial density exacerbates LATE-NC–related hippocampal atrophy independent of AD pathologic diagnosis.

## DISCUSSION

In this study linking ex-vivo MRI and neuropathology, higher microglia density was associated with lower hippocampal-to-hemisphere volume ratio. However, this association was substantially attenuated after accounting for multiple coexisting neurodegenerative and cerebrovascular pathologies. Furthermore, we find that the association of LATE-NC, but not AD pathology, with hippocampal-to-hemisphere volume ratio is stronger in the context of greater microglia density.

Microglia are the primary immune cells of the central nervous system and are highly responsive to age-related neuronal stress. In our prior work, we demonstrated that microglia inflammation in the hippocampus is an important associate of cognitive decline and that both AD pathology, specifically tangles, and LATE-NC are independently associated with regionally elevated microglia density within the hippocampus ^14^. Findings from the current study extend upon the complex relationship between LATE-NC, AD, and microglial inflammation on neurodegenerative processes. Our findings suggest that the relationship between microglia and hippocampal atrophy is not an independent mechanism in hippocampal volume loss, and that hippocampal atrophy is largely associated with AD and LATE brain pathologies. Indeed, β-amyloid burden, tangle density, and LATE-NC each demonstrated independent associations with smaller hippocampal-normalized volumes, consistent with prior studies implicating these lesions in medial temporal lobe vulnerability ^27–30^. Further, we identified a significant interaction between LATE-NC and microglial density, indicating a synergistic relationship on hippocampal volume. Higher microglial density exacerbated the deleterious association of LATE-NC with hippocampal volume. Interestingly, this pattern differed from AD pathology, in which we did not find an interaction between microglia with β-amyloid or tangles. Instead, regardless of microglia density, β-amyloid and tangles remained robustly associated with lower hippocampal-to-hemisphere volume ratio. These data suggest that AD-related hippocampal atrophy may proceed through biological pathways that are not strongly dependent on, or influenced by, microglial density within the hippocampus. Whereas hippocampal microglia inflammation may play a more central role in the hippocampal vulnerability associated with LATE-NC.

Hippocampal atrophy remains a key biomarker of neurodegeneration in the NIA-AA research framework for staging recently revised AD staging criteria ^31^. However, hippocampal atrophy is not specific to AD and can arise from multiple underlying pathologic processes. LATE-NC is defined by the accumulation of cytoplasmic TDP-43 aggregates across limbic (amygdala and hippocampus) and mesial temporal cortices, with extension into the frontal lobe in advanced stages ^32^. We and others have shown that LATE-NC is strongly associated with disproportionate hippocampal atrophy, often exceeding the degree of structural loss expected from AD pathology alone ^4,29,33,34^. Mechanistic studies have shown that dysregulation of TDP-43 expression has been associated with increased microglial activation and innate immune signaling ^35^. While this response may initially be adaptive in the brain environment (i.e., removing dysfunctional synaptic elements or clearing cellular debris), prolonged chronic activation can become maladaptive, shifting microglia cells to a pro-inflammatory state. Furthermore, the accumulation of senescent microglia with age may lead to the acceleration of brain aging and increased vulnerability to neurodegenerative processes ^36,37^. In the current study, our findings from stratified analyses further suggested that microglial burden exacerbates LATE-NC–related hippocampal atrophy independent of AD pathologic diagnosis.

The lack of interaction between microglia and AD pathologic markers on hippocampal-to-hemisphere volume ratio was intriguing, suggesting that microglia within the hippocampus have limited influence on hippocampal atrophy attributable to amyloid or tau. It is also possible that because the association between microglia and LATE-NC is so strong that the effects of AD, particularly tangle pathology, on hippocampal volume loss become difficult to separate. It is well described that hippocampal atrophy is a prominent neuroimaging feature observed in AD. Our findings certainly point to perhaps microglia-independent mechanisms in which AD pathology contributes to hippocampal vulnerability. Notably, synaptic dysfunction and disrupted connectivity between hippocampal and other mesial temporal lobe brain regions, tau seeding, and microtubule destabilization and impaired axonal transport are some of the plausible mechanisms in which AD may impact structural vulnerability of the hippocampus ^38,39^.

This study offers new insight into the complex interplay between neurodegenerative proteinopathies, microglia inflammation, and structural MRI changes in the hippocampus, which may have important mechanistic and therapeutic implications. An important strength of this study is the utilization of digital pathology to obtain quantitative measures of microglia pathology, capturing the full range of microglia burden in the hippocampus. Additionally, this study included a wide variety of age-related neuropathologies, including microglial inflammation, to examine contributions to hippocampal atrophy. A noteworthy limitation to this study is that all MRI and neuropathologic assessments were performed postmortem, preventing inferences about the temporal sequence of microglial activation, LATE-NC progression, and hippocampal atrophy. A single marker was used to quantify microglia, and while CR3-43 labels a subset of microglia/macrophages, other microglia markers may be differentially related to neurodegeneration. Although we adjusted for multiple age-related neuropathologies, residual confounders from unmeasured processes cannot be excluded. Lastly, participants were predominantly non-Hispanic White, very old, and largely female, which raises questions to be addressed in other populations.

## Acknowledgements

We thank all the participants from the Rush Memory and Aging Project, Religious Orders Study, and Minority Aging Research Study, the RADC clinical Core, and Latino Core Study for enrollment and brain donation. We also want to thank investigators and key staff members at Rush Alzheimer’s Disease Center; Traci Colvin, Tracey Nowakowski and Shayla Calloway, for study coordination; and Srabani Mondal and Ryan Johnson for neuropathology lab supervision; John Gibbons and Greg Klein for data management.

## Funding

This work was supported by National Institute of Health grants: R01AG017917, R01AG067482, P30AG072975, R01AG022018, R01AG064233, U01NS100599, R01AG052200, RF1NS139975, and K01AG075177

## Author Contributions

Dr Kapasi trained and optimized the pathology classifier for hippocampal microglia, and Dr. Chen ran the image analysis on hippocampal sections. Drs. Kapasi and Leurgans were involved in validating the pathology classifier. Drs. Kapasi, Leurgans, and Schneider were involved in the conceptional, organization, and execution of the project. Drs. Leurgans and Yu were involved in the review of the statistical analyses. Dr. Arfanakis was involved in the review of the ex-vivo MRI data. Dr. Kapasi wrote the first draft, and Drs. Yu, Leurgans, Agarwal, Bennett. Arfanakis, and Schneider reviewed and critiqued manuscript drafts for intellectual content.

## Disclosures

All authors have no competing interests to declare.

## Notes

### Competing Interest Statement

The authors have declared no competing interest.

