## Supplementary Table 1 for "Differential Associations of Microglial Inflammation on LATE-NC and Tangle-Related Hippocampal Atrophy"

**Supplementary Table 1.** Interaction of AD pathology and LATE-NC with stage 2/3 microglia density in relation to hippocampal volume**.**

| **Predictors** | **Interaction** | **Hippocampal to Hemisphere Volume Ratio**  **Estimate (SE, P-value)** |
| --- | --- | --- |
| β-amyloid |  | -2.886 (1.17, 0.01) |
| Stage 2/3 Microglia |  | 0.386 (3.69, 0.92) |
|  | β-amyloid x Stage 2/3 Microglia | 0.084 (2.47,0.97) |
| Tangles |  | -1.928 (0.90,0.03) |
| Stage 2/3Microglia |  | -3.459 (4.11,0.40) |
|  | Tangle x Stage 2/3 Microglia | 2.455 (1.80,0.17) |
| LATE-NC |  | -1.475 (0.858,0.08) |
| Stage 2/3 Microglia |  | 6.798 (3.21,0.04) |
|  | LATE-NC x Microglia | -3.914 (1.54,0.01) |

Estimates from linear regression models in which hippocampal-to-hemisphere volume ratio was the outcome. Models included an interaction between stage2/3 microglia with either β-amyloid, tau tangle burden, or LATE-NC, and adjusted for demographics, scanner, PMI to fixation and imaging, and other age-related pathologies. In analyses, stage2/3 microglia data was log-transformed.
